# Larvicidal effect of some traditional Saudi Arabian herbs against *Aedes aegypti* larvae, a vector of dengue fever

**DOI:** 10.1101/2023.09.04.556245

**Authors:** Somia Eissa Sharawi

## Abstract

Mosquitoes are effective disease vectors for human and veterinary health because they share a close environment with humans and represent a major public health problem. Saudi Arabia is home to the endemic dengue fever disease, carried by the *Aedes aegypti* mosquito. Conventional insecticides based on organophosphates and insect growth regulators are the most effective short-term control methods for this vector. However, these insecticides are toxic to non-target organisms, the environment, and humans. This study, eight plant extracts (*Ilex paraguariensis, Camellia sinensis, Cinnamomum zeylanicum, Elettaria cardamomum, Matricaria chamomilla, Allium sativum, Coffea arabica*, and *Piper nigrum*) were assayed as an insecticide against the 3^rd^ and 4^th^ larval stage of *Ae. aegypti* (L.), after 24 and 48 h of exposure in the duration between March and June 2023. After 48h, all plants extracted showed the highest mortality (100%), except for *C. arabica* which showed the lowest mortality rate (98.33%) at 30%. *I. paraguariensis* showed the greatest effectiveness with an LC_50_ =7.17 ppm, followed by *P. nigrum* with an LC_50_ = 5.66 ppm. Further investigation is needed to purify the active ingredients responsible for their toxicity to mosquito larvae and to investigate the mechanisms of action of plant extracts in water and other solvents.

## Introduction

Among all insects in the order Diptera, mosquitoes (family Culicidae) are an effective disease vector for human and veterinary health **(Guzman et al., 2010)**, because they share a close environment with humans and represent a major public health problem. It is estimated that mosquitoes transmit diseases to more than 700 million people each year, and they are responsible for one in 17 deaths **(WHO, 2005)**. Saudi Arabia is home to the endemic dengue fever disease, carried by the *Aedes aegypti* mosquito **(Aziz et al., 2014)**. Dengue prevalence is increasing in the Middle East based on recent reports **(Aziz et al., 2014)**, showing a total of 4,411 dengue cases reported in this area. This high number of cases is associated with a remarkable failure of recent control strategies. Conventional insecticides are the most effective short-term control measure against this vector **(Malik et al., 2007)**. Most of them are based on organophosphates and insect growth regulators **(Yang et al., 2002)**. These applications have shown a knockout effect during outbreaks, and several environmental and health issues have emerged during insecticide application **(Rahuman et al., 2008)**. In addition to toxicity to non-target organisms, the effects on the environment and human health have also increased **(Lee et al., 2001)**. In contrast, similar to other insects, mosquitoes develop resistance to insecticides following their application. Further research is needed to find new natural insecticides that can reduce the impact of chemicals on the environment and human health. Plants and their derivative compounds are an appropriate alternative because of their effectiveness and lower toxicity to human health and the environment than conventional insecticides. Numerous techniques can be used to extract plant parts for industrial and commercial use in liquid or solid forms. Maceration is one of the most widely used and affordable methods to extract various bioactive plant chemicals, which involves steeping coarse and powdered plant materials in various solvents **(Saqib et al., 2022)**. In recent decades, many studies have shown the efficiency of different plant extracts, such as mosquito larvicides, repellents, over deterrents, and growth inhibitors, without causing any harm to humans **(Saqib et al., 2022)**. The mechanisms of action of plant extracts on mosquito larvae include neurotoxic effects, inhibition of detoxifying enzymes, disruption of larval growth, and/or injury to the midgut **(Pavela et al., 2019)**. According to **(Pavela et al., 2019)**, 29 of 400 examined plant species showed outstanding larvicidal activity against *Aedes* species. **Ramanibai et al., 2016** reported that *Annona squamosa* extracts elicited toxicity against all stages of *Ae. aegypti* after 24 h. When used against first-instar *Aedes* larvae, an aqueous *Lantana camara* leaf extract achieved the highest mortality rate at a concentration of 400 mg/100 mL concentration after 96 h **(Sivakumar et al., 2022). Elkhidr et al., 2020**, observed mortality rates between 75 and 100% for the aqueous extract of *Acacia nilotica* on *Culex* species larvae in laboratory conditions using successive concentrations of 0.0125–2%. Numerous studies support the use of plant extracts and their derivatives as a safe and effective measure to control mosquito larval stages; hence, this study aimed to examine the insecticide effects of macerated extracts of mate (*Ilex paraguariensis*), green tea (*Camellia sinensis*), cinnamon (*Cinnamomum zeylanicum*), cardamom (*Elettaria cardamomum*), chamomile (*Matricaria chamomilla*), garlic (*Allium sativum*), coffee (*Coffea arabica*), and black pepper (*Piper nigrum*) as biological controls against the 3^rd^ and 4^th^ stages of *Ae. aegypti*.

## Materials and methods

### Mosquitoes culture

An experimental investigation was conducted to determine the toxic effects of mate (*I. paraguariensis*), green tea (*C. sinensis*), cinnamon (*C. zeylanicum*), cardamom (*E. cardamomum*), chamomile (*M. chamomilla*), garlic (*A. sativum*), coffee (*C. arabica*), and black pepper (*P. nigrum*) macerated extracts on 3^rd^ and 4^th^ stage *Ae. aegypti* larvae. Experiments were conducted at the dengue mosquito research station (Saudi Arabia) at King Abdelaziz University in Jeddah in the duration between March and June 2023. This study used *Ae. aegypti* laboratory strains, and the laboratory conditions were 27 ± 1 °C, 70 ± 5% relative humidity, and 14:10 (L:D). Fish food or dry bread powder mixed 1:1 with dried milk was fed to the larvae. For quality control, all collected mosquito larvae were checked for species and viability by an experienced enterologist.

### Plant extraction

The plants were purchased from local markets. Mate (*I. paraguariensis*), green tea (*C. sinensis*), cinnamon (*C. zeylanicum*), cardamom (*E. cardamomum*), chamomile (*M. chamomilla*), garlic (*A. sativum*), coffee (*C. arabica*), and black pepper (*P. nigrum*) were washed, dried in a shaded area, powdered using a clean electric blender, and prepared for extraction. Twenty grams of each plant were soaked in 100 mL of distilled water for 24 h at room temperature (37 °C). Further serial dilutions were done from the above stock solutions (1, 5, 10, 20, and 30%).

### Mortality bioassay

Each plant stock solution was created by mixing 1 mL of the solution with 99 mL of distilled water. Twenty larvae were placed in small cups containing 50 mL of each concentration. In addition, 20 larvae were placed in a separate cup containing 50 mL of distilled water as a control. Mortality was assessed after 24 and 48 h by counting the dead and live insects. Larvae were considered dead when they settled at the bottom without moving. Experiments were conducted at room temperature (30-37 °C). Each experiment was repeated thrice.

### Statistical analysis

This investigation was conducted using a random design. Mortality in the treatment groups was corrected using Abbott’s formula if the mortality of the control was between 5 and 20% (Abbott, 1925). The LC_50_ and LC_90_ were calculated using the Probit analysis program. Larval mortality was recorded daily.

## Results and discussion

Our findings demonstrate a connection between concentration and mortality, which was either modest or moderate. The larval mortality rates were substantially higher in the treated groups than in the control group. The data presented in Tables 1-8 show the effects of different plant extract concentrations on the 3^rd^ and 4^th^ larval instars of *Ae. aegypti* after 24 and 48 hours of exposure. After 24 h, *E. cardamomum* and *M. chamomilla* showed the highest mortality rates (60%), followed by *I. paraguariensis, C. zeylanicum, C. arabica*, and *P. nigrum* (58%). *C. sinensis* had the lowest mortality rate with (54%) after 24 h. After 48 h at 10%, *C. zeylanicum* showed the highest mortality (96.7%) against larvae, followed by *M. chamomilla* (88.33%), *A. sativum* (81.67%), *C. sinensis* (75%), *C. arabica* (71.67%), *P. nigrum* (68.33%), and *E. cardamomum* (66.67%). *I. Paraguariensis* caused the lowest mortality rate after 48 h (48.33%). The leaves of *I. paraguariensis* and *M. chamomilla* are commonly used in tea infusions in the Middle East. However, little research has been conducted on the larvicidal effectiveness of their aqueous extracts against *Ae. Aegypti* larvae. Previous studies on health benefits have revealed antimutagenic, antioxidant, hepatoprotective, and glycemic improvements. In our study, the mean mortality of *I. paraguariensis* and *M. chamomilla* was 20% after 48 h. To date, there are no studies to confirm our results. A study on *C. sinensis* confirmed our results, showing that the exposure of larval-stage *Anopheles gambiae* to a crude *C. sinensis* extract at 250 ppm and 500 ppm for 24 h caused larval mortality rates of over 90%, as well as 75% in *Anopheles arabiensis* **(Muema et al., 2016)**. A relatively lower concentration of 100 ppm resulted in moderate mortality rates of <50% in both species. For *C. zeylanicum* and *E. cardamomum* (Tables 3 and 4), 100% mortality was observed after 48 h of exposure. According to **(Sanei-Dehkordi et al., 2022)**, *C. zeylanicum* and *E. cardamom* essential oils caused 100% larval death against *Anopheles stephensi*. ^**[15]**^ noted that *A. sativum* extract exposure time and extract concentration increased *Culex* and *Anopheles* mortality. The highest mortality was observed at 3.0 mg/mL after 3 h of exposure for both *Culex* and *Anopheles* at a mean of 10.00 each, while 0.5 mg/mL recorded the lowest mortality after 1 h of exposure with means of 2.33 and 3.67, respectively. These results are similar to our study, where 30% caused 100% mortality after 48 h (Table 6). *C. arabica* caused 58% and 98% mortality after 34 and 48 h of exposure, respectively. Consistent with our findings, **(Wiwit et al., 2019)**, showed the insecticidal activity against *Ae. aegypti* of arabica coffee grounds. As seen in Table 8, *P. nigrum* showed the highest concentration of 30% after 48 h. Consistent with our results, the essential oils of *P. nigrum* were most effective against *A. gambiae* larvae, showing 100% mortality with an LC_50_ value of 149 ppm. As presented in Table 9 and Fig. 1, *C. zeylanicum* was the most effective plant extract after 48 h according to LC_50_, followed by *M. chamomilla, E. cardamomum, A. sativum, C. arabica, C. sinensis, P. nigrum*, and *I. paraguariensis* (1.12%, 2.27%, 3.58%, 3.61%, 4.34%, 4.50%, 5.66%, and 7.14%, respectively).

**Table (1):**
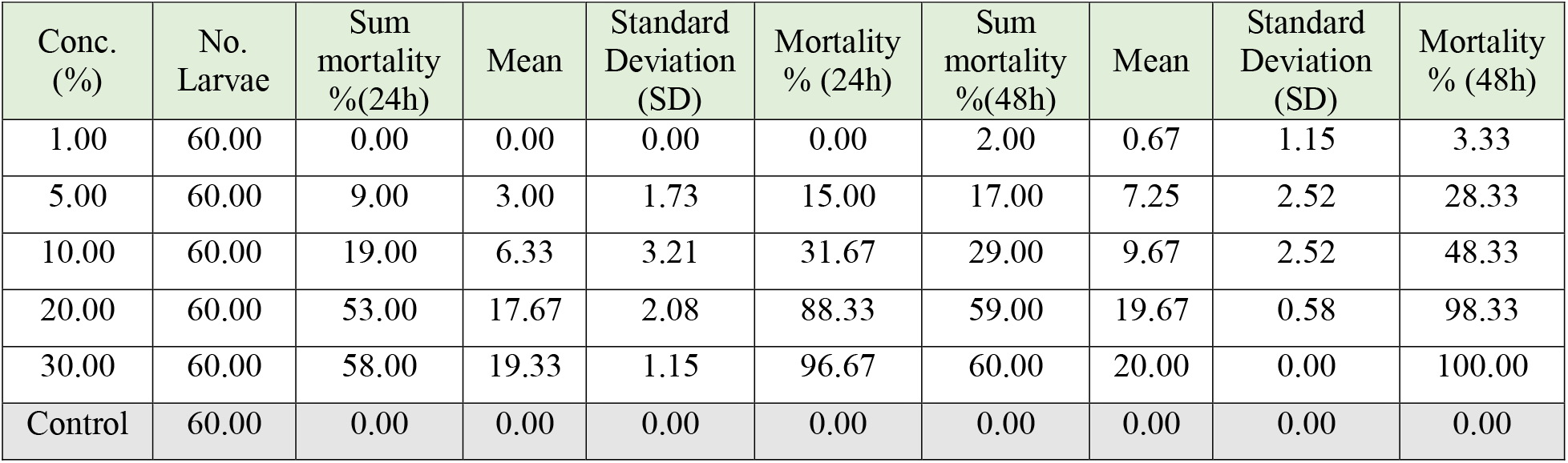
Effect of *I. paraguariensis* against *Ae. aegypti* larval stage after 24 and 48h.

**Table (2):**
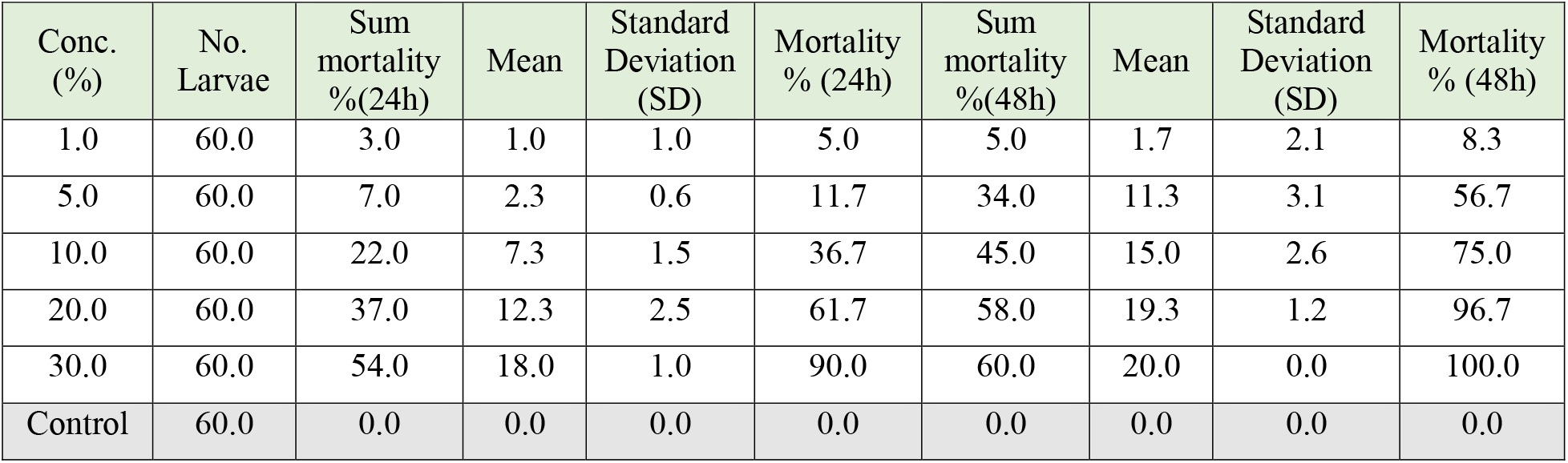
Effect of *C. sinensis* against *Ae. aegypti* larval stage after 24 and 48h.

**Table (3):**
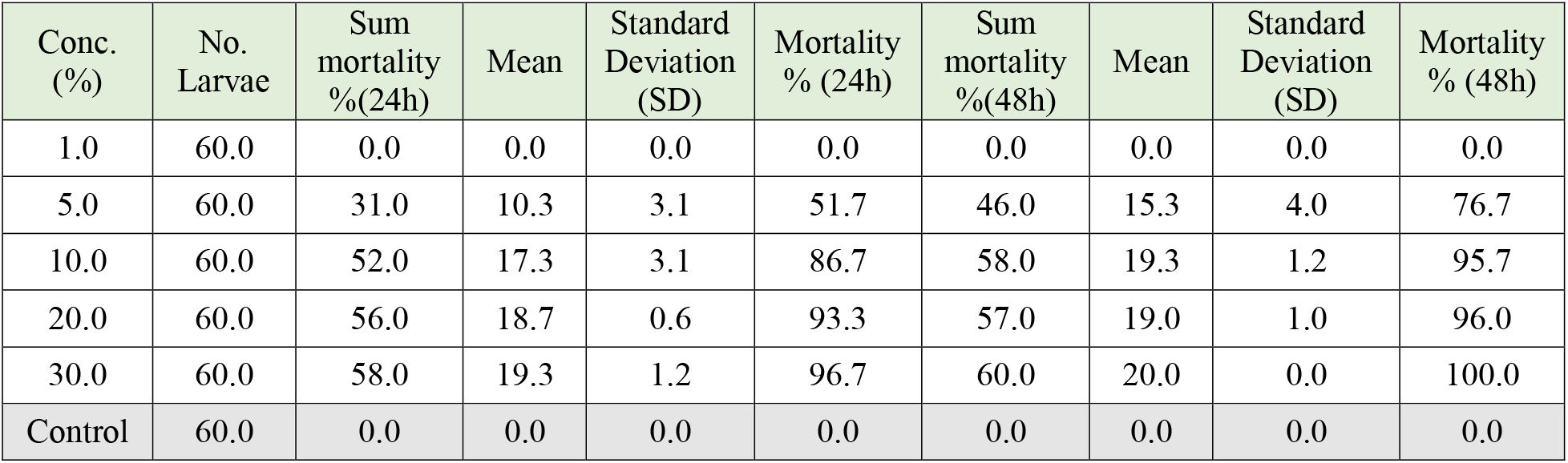
Effect of *C. Zeylanicum* against *Ae. aegypti* larval stage after 24 and 48h.

**Table (4):**
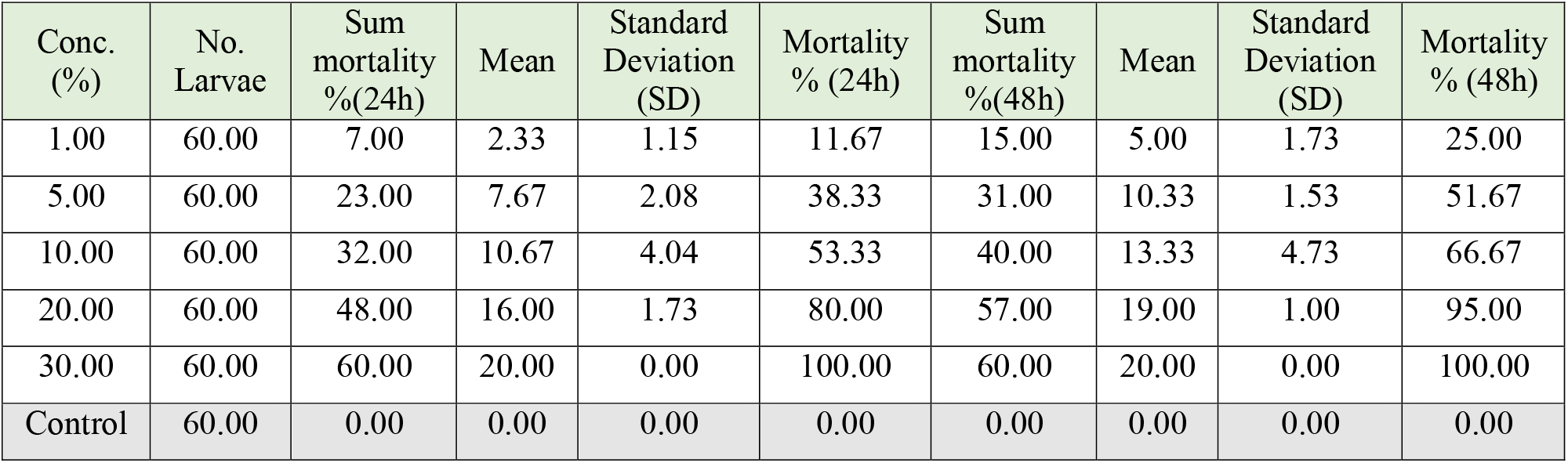
Effect of *E. cardamomum* against *Ae. aegypti* larval stage after 24 and 48h.

**Table (5):**
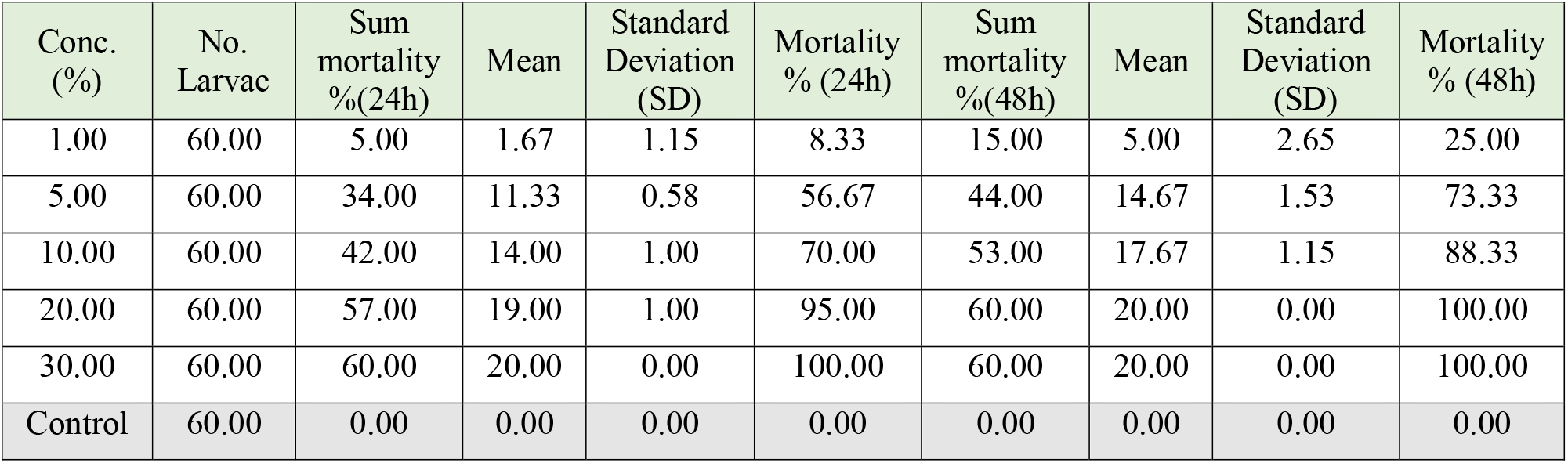
Effect of *M. chamomilla* against *Ae. aegypti* larval stage after 24 and 48h.

**Table (6):**
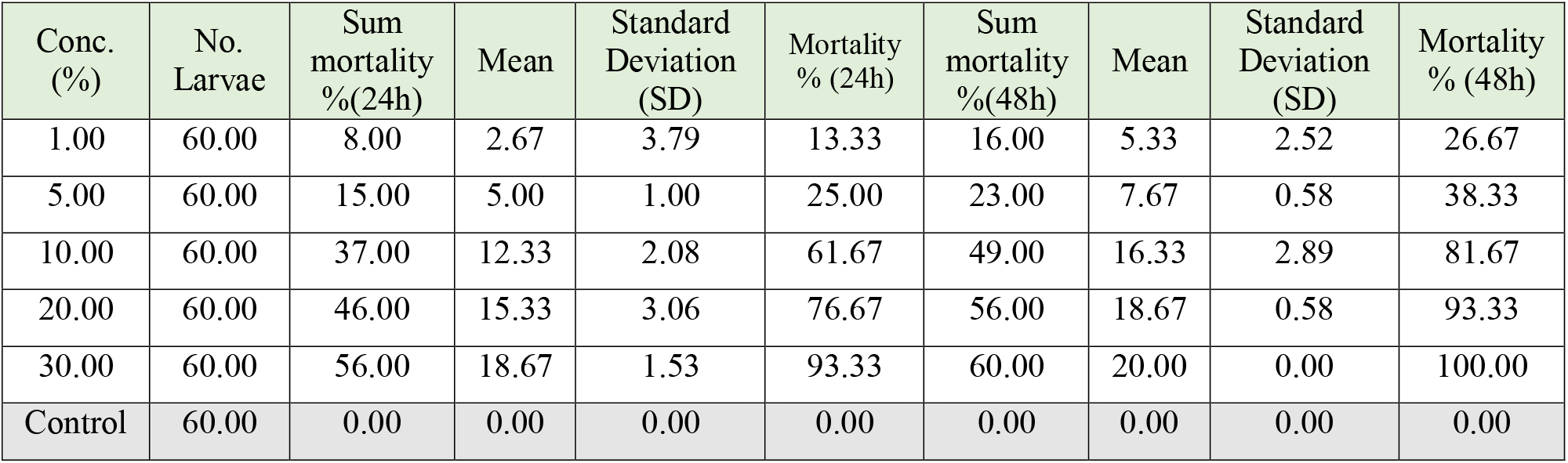
Effect of *A. sativum* against *Ae. aegypti* larval stage after 24 and 48h.

**Table (7):**
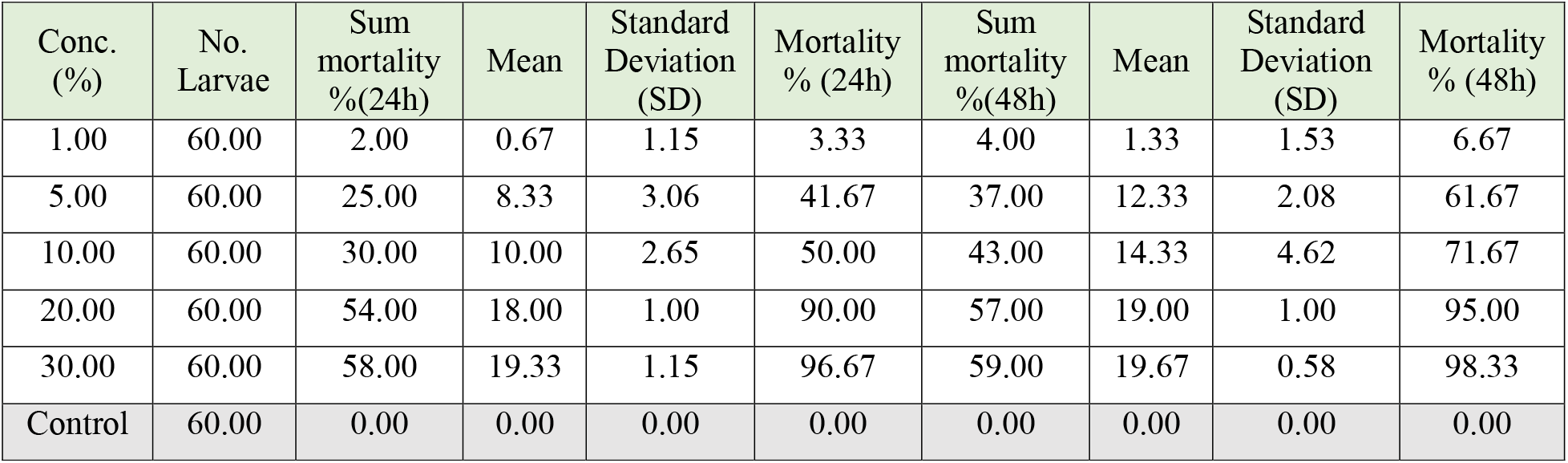
Effect of *C. arabica* against *Ae. aegypti* larval stage after 24 and 48h.

| Conc. (%) | No. Larvae | Sum mortality % (24h) | Mean | Standard Deviation (SD) | Mortality % (24h) | Sum mortality % (48h) | Mean | Standard Deviation (SD) | Mortality % (48h) |
| --- | --- | --- | --- | --- | --- | --- | --- | --- | --- |
| 1.00 | 60.00 | 2.00 | 0.67 | 1.15 | 3.33 | 4.00 | 1.33 | 1.53 | 6.67 |
| 5.00 | 60.00 | 25.00 | 8.33 | 3.06 | 41.67 | 37.00 | 12.33 | 2.08 | 61.67 |
| 10.00 | 60.00 | 30.00 | 10.00 | 2.65 | 50.00 | 43.00 | 14.33 | 4.62 | 71.67 |
| 20.00 | 60.00 | 54.00 | 18.00 | 1.00 | 90.00 | 57.00 | 19.00 | 1.00 | 95.00 |
| 30.00 | 60.00 | 58.00 | 19.33 | 1.15 | 96.67 | 59.00 | 19.67 | 0.58 | 98.33 |
| Control | 60.00 | 0.00 | 0.00 | 0.00 | 0.00 | 0.00 | 0.00 | 0.00 | 0.00 |

**Table (8):** Effect of *P. nigrum* against *Ae. aegypti* larval stage after 24 and 48h.

| Conc. (%) | No. Larvae | Sum mortality % (24h) | Mean | Standard Deviation (SD) | Mortality % (24h) | Sum mortality % (48h) | Mean | Standard Deviation (SD) | Mortality % (48h) |
| --- | --- | --- | --- | --- | --- | --- | --- | --- | --- |
| 1.00 | 60.00 | 1.00 | 0.33 | 0.58 | 1.67 | 4.00 | 1.33 | 1.53 | 6.67 |
| 5.00 | 60.00 | 17.00 | 5.67 | 1.15 | 28.33 | 23.00 | 7.67 | 2.08 | 38.33 |
| 10.00 | 60.00 | 32.00 | 10.67 | 1.15 | 53.33 | 41.00 | 13.67 | 0.58 | 68.33 |
| 20.00 | 60.00 | 52.00 | 17.33 | 0.58 | 86.67 | 55.00 | 18.33 | 0.58 | 91.67 |
| 30.00 | 60.00 | 58.00 | 19.33 | 0.58 | 96.67 | 60.00 | 20.00 | 0.00 | 100.00 |
| Control | 60.00 | 0.00 | 0.00 | 0.00 | 0.00 | 0.00 | 0.00 | 0.00 | 0.00 |

**Table (9):** LC_50_ and LC_90_ of plant extracts against *Ae. aegypti* larval stage after 48h

|  | LC<sub>50</sub> | LC<sub>90</sub> | Slope |
| --- | --- | --- | --- |
| *I. paraguariensis* | 7.14 | 17.86 | 3.22 |
| *C. sinensis* | 4.50 | 18.44 | 2.09 |
| *C. Zeylanicum* | 1.12 | 7.90 | 1.51 |
| *E. cardamomum* | 3.58 | 25.62 | 1.51 |
| *M. chamomilla* | 2.27 | 9.55 | 2.05 |
| *A. sativum* | 3.61 | 26.68 | 1.47 |
| *C. arabica* | 4.34 | 15.73 | 2.29 |
| *P. nigrum* | 5.66 | 19.12 | 2.42 |

**Fig. (1)):**
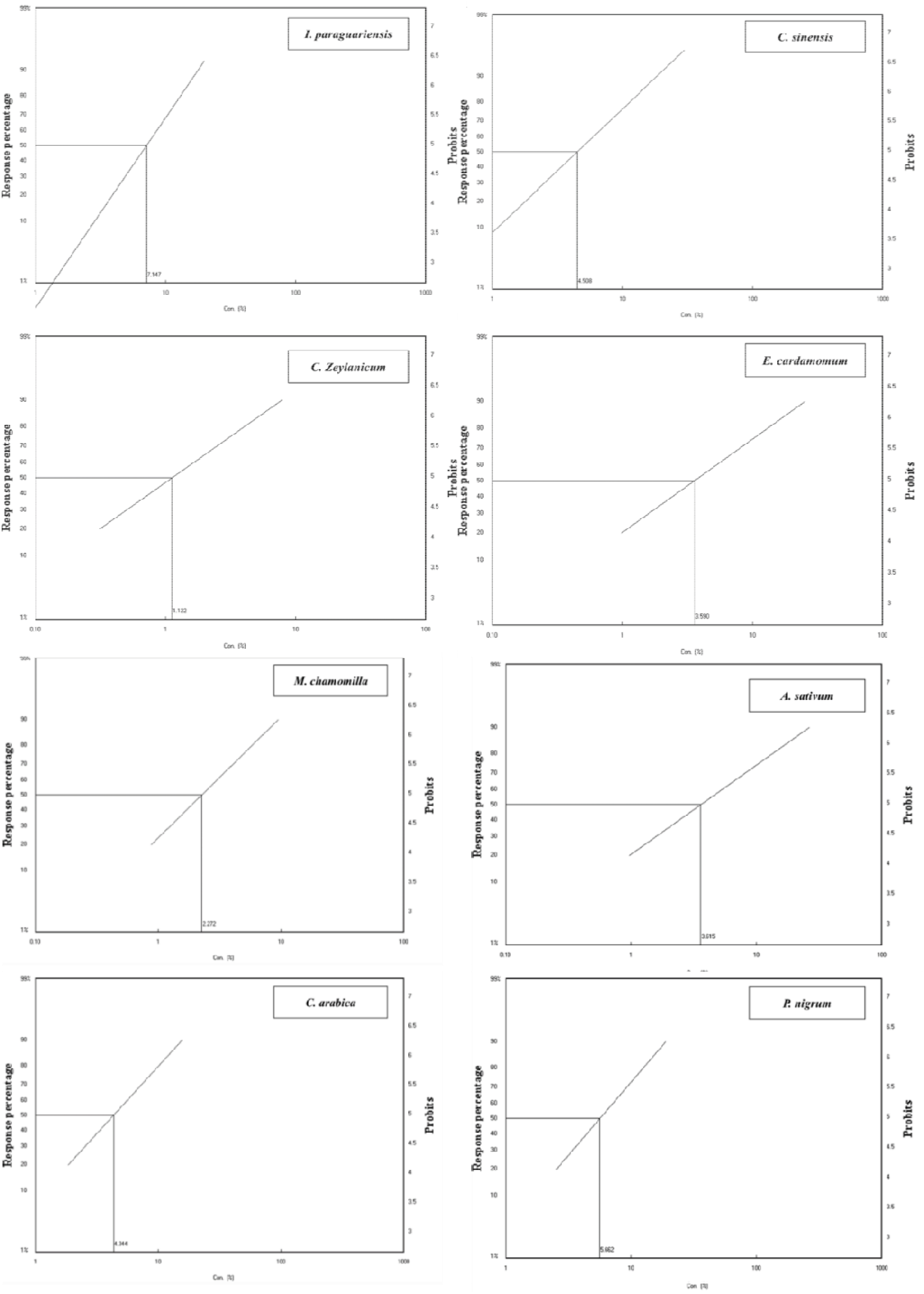
LC_50_ of plant extracts against *Ae. aegypti* larval stage after 48h.

## Conclusion

This study aimed to examine the insecticidal effects of macerated extracts of *I. paraguariensis, C. sinensis, C. zeylanicum, E. cardamomum, M. chamomilla, A. sativum, C. arabica*, and *P. nigrum* against 3^rd^ and 4^th^ stage *Ae. aegypti* larvae under laboratory conditions after 24 and 48 h of exposure. *C. zeylanicum* was the most effective plant extract, while *I. paraguariensis* caused the lowest toxicity. These plant extracts may provide additional methods to reduce *Ae. aegypti* populations. These findings might motivate researchers to evaluate novel, naturally occurring, and active chemicals that are safer and more manageable than manufactured insecticides.

## Author contribution

Dr. Somia Sharawi conceived and designed the study conducted research, provided research materials, and collected and organized data. Also, analyzed and interpreted data, wrote the initial and final draft of the article, and provided logistic support. The author has critically reviewed and approved the final draft and is responsible for the content and similarity index of the manuscript

## Ethical approval

There is no ethical issue

